# A single domain intrabody targeting the follicle-stimulating hormone receptor (FSHR) impacts FSH-induced G protein-dependent signalling

**DOI:** 10.1101/2023.08.07.552286

**Authors:** Pauline Raynaud, Vinesh Jugnarain, Océane Vaugrente, Amandine Vallet, Thomas Boulo, Camille Gauthier, Asuka Inoue, Nathalie Sibille, Christophe Gauthier, Frédéric Jean-Alphonse, Eric Reiter, Pascale Crépieux, Gilles Bruneau

## Abstract

Intracellular variable fragments from heavy-chain antibody from camelids (intra-VHH) have been successfully used as chaperones to solve the 3D structure of active G protein-coupled receptors bound to their transducers. However, their effect on signalling has been poorly explored, although they may provide a better understanding on the relationships between receptor conformation and activity. Here, we isolated and characterized iPRC1, the first intra-VHH recognizing a member of the large glycoprotein hormone receptors family, the follicle-stimulating hormone receptor (FSHR). This intra-VHH recognizes the FSHR 3^rd^ intracellular loop and decreases cAMP production in response to FSH, without altering Gαs recruitment. Hence, iPRC1 behaves as an allosteric modulator and provides a new tool to complete structure/activity studies performed so far on this receptor.

## Introduction

G protein-coupled receptors (GPCRs) represent a large family of membrane receptors characterized by an N-terminal extracellular domain, seven transmembrane domains connected by three extracellular and three intracellular loops, and an intracellular C-terminal domain. Ligand binding to a GPCR leads to conformational changes within the transmembrane and intracellular domains, mediating the transduction of an extracellular signal to the interior of the cell. GPCRs are highly dynamic proteins, adopting a plethora of active and inactive conformations (1). This is in accordance with the fact that, upon ligand binding, they are able to interact with a diversity of downstream signalling proteins with variable efficacy. The signal is transduced by heterotrimeric G proteins (Gs, Gi/o, Gq/11, G12/13), G protein-coupled receptor kinases (GRK) and β-arrestins among others. Since GPCRs are involved in the regulation of numerous physiological and pathological processes, regulating their biological activity has been the object of considerable efforts, involving the use of a variety of molecular tools. Ligands recognizing the orthosteric binding site of a GPCR act as agonists or antagonists. The use of allosteric modulators that favour either active (positive allosteric modulator) or inactive (negative allosteric modulator) conformations may fine-tune GPCR signalling. Biased ligands, whether orthosteric or allosteric, exhibit altered efficacy or potency to regulate a specific signalling pathway, when compared to a reference ligand, providing valuable tools to study and control GPCR signalling regulation.

Small molecules have been extensively explored to modulate GPCR signalling and pharmacology, including the FSHR’s (2). In the last decade, there has been a growing interest in the development of antibodies targeting GPCRs, especially in the VHH (VH of heavy chain only antibodies) format. These VHHs of low molecular weight (∼13 kDa), display the smallest paratope capable of specifically recognizing an epitope. A long and flexible CDR3 that compensates for the absence of light chain enables them to access and recognize cryptic epitopes, such as binding pockets (3). Furthermore, their conformational stability is an advantageous property. VHHs have disulphide bonds not essential for epitope recognition (4), hence they are especially well-suited for intracellular expression despite reducing conditions in the cytosol.

Ligand-bound GPCRs are structurally instable, and the recruitment of transduction proteins (G proteins, β-arrestins) each stabilize a discrete subset of receptor conformations (5). Similarly, intracellular VHHs (intra-VHHs) have the ability to stabilize selective GPCR conformations. Therefore, intra-VHHs targeting GPCRs have been a tremendous help to solve crystal structures of numerous active GPCRs. For example, Nb80 was used as a chaperone to solve crystal structure of the active beta 2 adrenergic receptor (ADRB2) (6), Nb9-8 for the structure of active muscarinic acetylcholine receptor M2 (7), or Nb39 for the active structure of mu-opioid receptor (8). All of these intra-VHHs exhibit G protein mimetic properties. Intra-VHHs stabilising partially active conformations of GPCRs have also been identified, such as Nb71 targeting ADRB2 (9). Others were able to stabilize inactive conformations, as Nb60, which acts as a negative allosteric modulator at the ADRB2 (10). In addition, some intra-VHHs have been used as biosensors to track GPCR intracellular trafficking. For instance, Nb80 fused to GFP as a biosensor for ADRB2 activation, revealed that active ADRB2 is present at the endosomes (11). Overall, by providing important stabilizing tools, intra-VHHs allowed valuable structural insights on GPCR activation. However, their modulatory impact on GPCR functioning has been poorly explored. So far, few studies reported a functional allosteric effect of intra-VHHs on GPCR signalling, and they are all directed against five GPCRs only (OPRK1, OPRD1, OPEM1, ADRB2 and US28), amongst the 800 members of the family (for review see Raynaud et al., 2022 (12)). For example, eighteen intra-VHHs selected on ADRB2 have been functionally characterized and they display variable inhibitory effects on cAMP production and β-arrestin recruitment (13). Four others inhibited the cAMP-responsive element (14). Intra-VHHs used as regulators of GPCR function can provide insight on GPCR signalling and on the receptor regions that are involved.

The follicle-stimulating hormone receptor (FSHR) is a class A (rhodopsin-like) GPCR playing a key role in reproduction, by regulating gametogenesis in male and female (15). The FSHR, together with the luteinizing hormone/choriogonadotropin (LHCGR) and thyroid stimulating hormone receptors (TSHR), constitute the subfamily of glycoprotein hormone receptors. Unlike other class A GPCRs, these three receptors are characterized by a large horseshoe-shaped extra-cellular domain, comprising the hormone binding domain (16). The FSH-stimulated FSHR primarily signals through Gs, the Gαs subunit leading to the production of the cAMP second messenger, thus activating protein kinase A (PKA). The Gβγ subunits activate GRKs, which subsequently phosphorylate the receptor on specific motifs, leading notably to the recruitment of β-arrestins (17–19). The latter play a key role in receptor desensitization, internalization and recycling, but also in G protein-independent signalling. FSH-activated FSHR is also engaged with other interaction partners (20), and induces a complex signalling network (21).

Through its biological role, the FSHR is a potential target for non-hormonal alternatives to fertility treatments or contraceptives, with the aim of optimizing efficacy while reducing side effects. However, it is not yet clearly understood how a specific signalling pathway correlates with the induced physiological effect. Therefore, to be able to fine-tune FSHR signalling, there is a need for tools allowing to decipher the complexity of the FSHR signalling network. In this study, we isolated a VHH targeting the 3^rd^ intracellular loop of the FSHR, expressed it as an intra-VHH in mammalian cells, showed its binding to the FSHR, and explored its potential modulatory effect on the FSH-induced Gs signalling pathway.

## Materials and methods

References of the materials are provided in Supplementary Table 1. Commonly used methods i.e. cell imaging, flow cytometry, VHH production and purification, BRET assays and cAMP measurement, are described in the supplementary methods section.

### Phage library

The phage library was obtained after intramuscular immunisation of llama with a cDNA encoding the human FSHR (In-Cell-Art, Nantes, France). Total RNAs were extracted from llama circulating leukocytes using LeukoLOCK™ Fractionation & Stabilization Kit (Invitrogen, Waltham, Massachusetts, USA) and reverse-transcribed as cDNA using SuperScript^TM^ III Reverse Transcriptase (Thermo Fisher Scientific, Waltham, Massachusetts, USA). VHHs cDNAs were amplified by nested PCR using Platinum^TM^ Taq DNA Polymerase (Thermo Fisher Scientific), as described in Supplementary Figure 1. PCR products were inserted between the SfiI and NotI sites of the pCANTAB 6 phagemid vector (kindly provided by Pierre Martineau). The resulting recombinant phagemids were transformed in *E. coli* TG1 bacteria (Lucigen, Teddington, UK). Addition of the KM13 helper phage (kindly provided by Pierre Martineau) allowed the production of the phages. The primary library contains 3.10^8^ independent phages.

### Phage display

Intra-VHH selection was performed by phage display on each of the three peptides corresponding to the intracellular loops of the human FSHR (ICL1, ICL2 and ICL3). ICL1 (TTSQYKLTVPR), ICL2 (ERWHTITHAMQLDCKVQLRH) and ICL3 (HIYLTVRNPNIVSSSSDTRIAKR) peptides were synthetized by Genecust (Boynes, France). Three selection rounds were carried out. For each round of selection, 1 µg of peptide in phosphate-buffered saline (PBS) pH 7.4 was coated into the selection well of Nunc MAXISORP strips (Dutscher, Bernolsheim, France). The first round started with a sample of 10^11^ to 10^12^ phages from the amplified phage library. Aspecific phages depletion and blocking of nonspecific sites were performed for 1 h on 2 wells coated with PBS 2 % bovine serum albumin (BSA) for round 1 and 3 and with PBS 2% skimmed milk for round 2. Phages were then incubated on the well coated with the peptides for 2 h. Phages were washed 40 times with PBS 0.1 % Tween 20, with one last wash in 50 mM Tris pH 8.0, 1 mM CaCl_2_. For rounds 1 and 2, phage elution was performed in the latter buffer containing 125 µg/mL bovine trypsin (Sigma-Aldrich, St. Louis, MO, USA), for 15 min at room temperature. For round 3, phages were eluted using 1.5 mM peptide in PBS pH 7.4. *E. coli* TG1 bacterial cells were used for phage amplification. Ninety-six randomly chosen clones among the selected clones from round 3 were sequenced (Azenta, Cambridge, MA, USA). Sequences were aligned, clustered and compared from one selection condition to the others. Sequences common to at least 2 selection conditions were considered as non-specific.

### Cell culture and plasmid transfections

Human Embryonic Kidney 293A (HEK293A) cells, HEK293/ΔGαs (a cell line depleted for Gαs protein) (22) and HEK293/ΔGαs/q/11/12/13 (23) were cultured in Dulbecco’s Modified Eagle Medium DMEM (Eurobio, Les Ulis, France) supplemented with 10% (v/v) foetal bovine serum, 100 IU/mL penicillin, 0.1 mg/mL streptomycin (Eurobio, Les Ulis, France) and kept at 37°C in a humidified 5% CO_2_ incubator. Cells were transiently transfected in suspension using the Metafectene Pro transfection reagent (Biontex Laboratories, München, Germany) following the manufacturer’s protocol. For interaction Bioluminescence resonance energy transfer (BRET) experiments, cells were transiently co-transfected with plasmids encoding the human FSHR C-terminally fused to the luciferase from the sea pansy *Renilla reniformis* (RLuc8) (0.1 μg plasmid DNA/cm^2^) as BRET donor, and increasing quantities of plasmids encoding the intra-VHH C-terminally fused to the Venus GFP derivative with a G4S linker as BRET acceptor (from 0 to 0.5 µg plasmid/cm^2^) (Supplementary figure 2A). To have an equivalent level of plasmid DNA transfected in all conditions, a pcDNA3.1 vector (Invitrogen, Waltham, Massachusetts, USA) encoding Venus alone was co-transfected. Further experiments were conducted with an excess of VHH plasmid DNA (0.45 µg DNA/cm^2^ of either T31-Venus or iPRC1-Venus plasmid) in addition to plasmids encoding human FSHR-RLuc8, human AVPR2-RLuc8, human AVPR1A-RLuc8, human AVPR1B-RLuc8, human CXCR4-RLuc8 or human PTH1R-RLuc8. Forty-eight hours following transfection, HEK293A cells were stimulated with either 3.3 nM FSH, 100 nM AVP, 250 nM SDF1α (Preprotech, Neuilly-sur-Seine, France) or 10 nM PTH (Bachem, Bubendiorf, Switzerland) according to the transiently expressed receptor, in presence of coelenterazine H. For Mini Gs protein (mGs, minimal engineered GTPase domain of the Gα subunit) recruitment BRET experiments, cells were transiently co-transfected with plasmids encoding the human FSHR-Rluc8 (0.1 μg plasmid DNA/cm^2^), the NES-Venus-mGs (65 ng plasmid DNA/cm^2^, kindly provided by Pr. Nevin A. Lambert, Augusta University, Augusta, GA, USA) (24,25), and VHH-6xHis (0.2 µg plasmid DNA/cm^2^) (Supplementary figure 2B). For HTRF and flow cytometry experiments, 0.5 ng Flag-tagged human FSHR (26) or 0.25 ng Flag-tagged human AVPR2 (27) plasmid DNA/cm^2^ and 80 ng of VHH-6xHis plasmid DNA/cm^2^ were used for transfection. For the condition without transfected VHH, an empty pcDNA3.1 plasmid was used. For confocal microscopy, two quantities of intra-VHH-Venus were tested for transfection (0.3 or 0.45 µg plasmid DNA/cm^2^). For Gαs protein competition, HEK293/ΔGαs/q/11/12/13 were transfected with 0.1 µg DNA/cm^2^ of FSHR-RLuc8 plasmid DNA, 0.25 µg DNA/cm^2^ of Venus-mGs or iPRC1-Venus plasmid DNA, and 0.5 µg DNA/cm^2^ of either empty pcDNA3.1 or Gαs protein plasmid DNA.

### Peptide competition by HTRF

Biotinylated peptides corresponding to FSHR ICL1, ICL2 and ICL3 (Genecust, Boynes, France) as previously defined, and non-biotinylated competing decapeptides ICL3-Nt (HIYLTVRNPN), ICL3-Mid (NPNIVSSSSD) and ICL3-Ct (SSSDTRIAKR) (GenScript Biotech, Piscataway, New Jersey, USA) were used. One hundred and twenty µM of a biotinylated peptide, with or without 1 mM of competing peptide were added to 30 nM of purified bacterially-expressed VHHs in PBS 0.1% Tween 20 (PBS-T), and incubated overnight at 4°C and 30 rpm. Background signal was obtained with PBS-T supplemented with equivalent amount of DMSO as the other conditions. The sensors, MAb Anti-6His-Tb cryptate (Cisbio, Waltham, Massachusetts, USA) and Streptavidin-d2 (Cisbio, Waltham, Massachusetts, USA) were added following manufacturer’s protocol, and after a 1-hour incubation in the dark, fluorescence measurement was performed with a TriStar² LB 942 Multimode Microplate Reader (Berthold Technologies GmbH & Co., Wildbad, Germany).

### In silico docking

The model of the iPRC1 VHH was generated by homology modelling using SWISS-MODEL (28–31). Docking of the iPRC1 CDR3 on the ICL3 of either inactive FSHR (PDB: 812H) or activated FSH-FHSR-Gs complex (PDB: 812G) was performed using the standard ‘EASY’ access settings on HADDOCK 2.4 (32,33). The generated HADDOCK models were visualised, superimposed onto each other, and analysed using the PyMOL Molecular Graphics System, Version 2.0 Schrödinger, LLC.

### Data analysis

Data were representative of at least 3 independent experiments and were expressed as mean values ± SD. Data were analyzed and plotted using Graphpad Prism 9 (Graphpad Software Inc., San Diego, CA, USA). For interaction BRET, nonspecific signal (when no VHH-Venus was expressed) was subtracted of the signal obtained with T31 or iPRC1, and data were then normalized to T31 and fitted to a specific binding curve with Hill slope (Graphpad Prism 9). For Mini Gs protein recruitment, BRET data were expressed as a percentage of the maximal FSH-induced response in the absence of VHH, and fitted to a one-phase association curve (Graphpad Prism 9). For HTRF cAMP accumulation assay, data were normalized to forskolin-induced cAMP accumulation and expressed as a percentage of ligand-induced response in the absence of intra-VHH. For competing peptides HTRF, signal was normalized to background and expressed as a percentage of iPRC1-ICL3 interaction signal. Statistical significance was estimated by the Mann-Whitney test to compare 2 different samples. Differences among means were considered significant at P < 0.05. Image processing was performed with Fiji (34).

### Data availability

The nucleotide sequence of iPRC1 has been deposited in Genbank with accession number OR554885

## Results

### Selection of the iPRC1 intra-VHH

To select intra-VHH targeting the FSHR intracellular side, we performed phage display on peptides corresponding to intracellular loops 1, 2 and 3 (ICL1, ICL2 and ICL3) of the receptor (Figure 1). Following three rounds of selection, the common clones identified in several conditions were considered as non-specific. Among the clones not common to ICL1 and ICL2 selections, a VHH directed against the FSHR ICL3, iPRC1, was selected through further analysis. An irrelevant intra-VHH, T31, randomly picked from a llama phage library and consistently observed as devoid of FSHR binding, has been used as a negative control in all following experiments.

**Figure 1:**
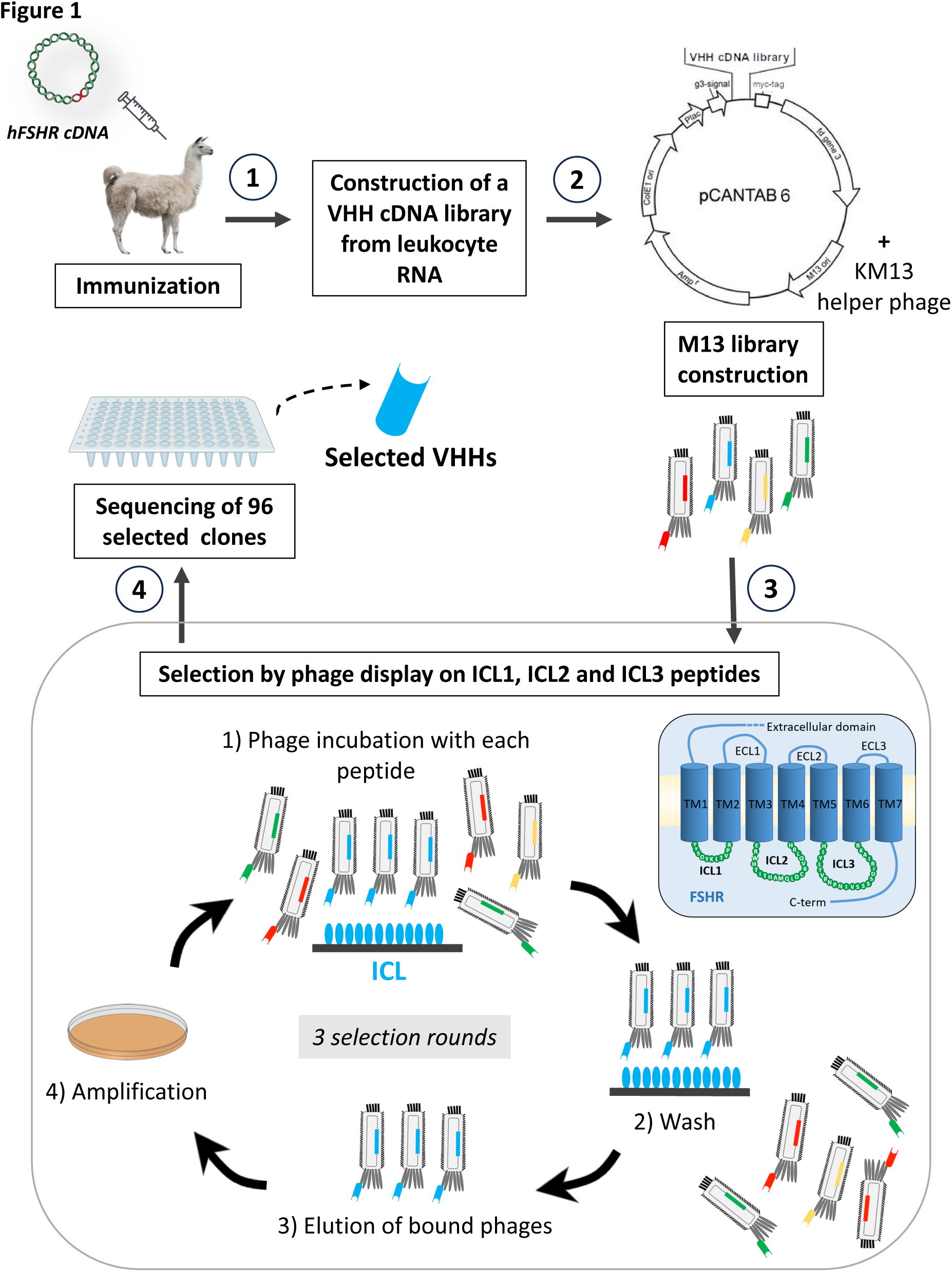
iPRC1 selection workflow. A phage library was constructed following llama immunization by injection of a cDNA encoding the hFSHR. Selection was done by phage display on the peptides corresponding to the FSHR intracellular loops, ICL1, ICL2 and ICL3. Ninety-six randomly selected clones were sequenced for each condition, and after cross-comparison, specific VHHs were selected.

### Expression of iPRC1 in mammalian cells

In order to be functionally characterized, iPRC1 and T31 were transiently expressed in HEK293A cells. We first assessed potential aggregation of the T31-Venus and iPRC1-Venus intra-VHHs (Figure 2A). Using confocal microscopy, we detected no sign of aggregation but rather homogeneous expression of both intra-VHHs, even with high quantities of transfected plasmid DNA (450 ng of DNA per cm^2^). Then the expression level of the two intra-VHH C-terminally linked to a 6xHis-tag was assessed by flow cytometry (Figure 2B), in all conditions of the following functional assays. Control and iPRC1 intra-VHHs both exhibited comparable levels of expression.

**Figure 2:**
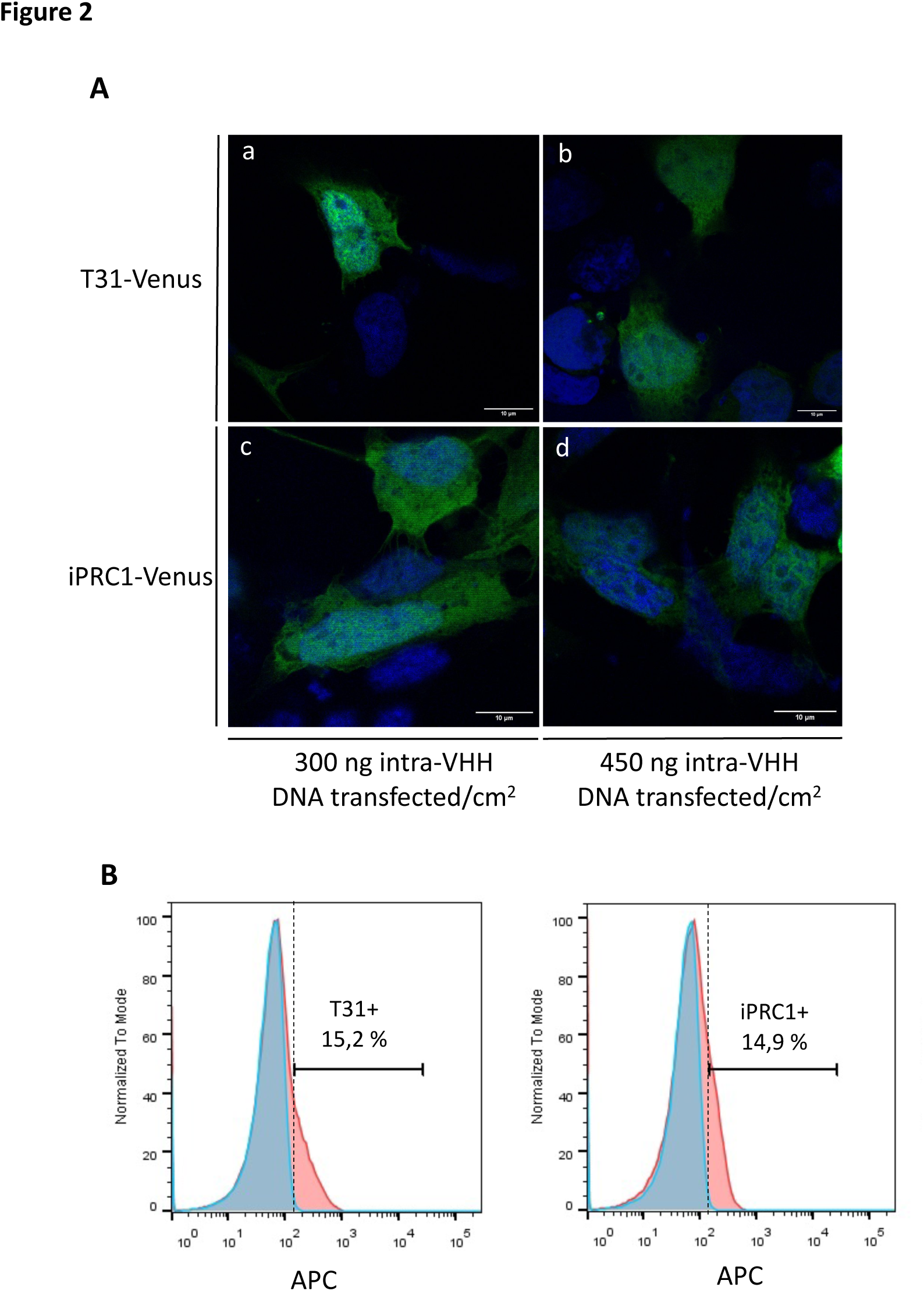
iPRC1 and irrelevant T31 intra-VHH expression in HEK293A cells. **(A)** Confocal microscopy representative images of T31-Venus (a and b) and iPRC1-Venus (c and d) (in green) transiently expressed in HEK293A cells, using two quantities of plasmid DNA (b and d). Cell nuclei were stained with Hoechst solution (in blue). **(B)** Flow cytometry histograms showing the population of cells expressing either the irrelevant intra-VHH T31-6xHis or iPRC1-6xHis (in red) in comparison to “mock”-cells transfected with empty pcDNA3.1 vector (in blue).

### iPRC1 binding to the FSHR

Even though iPRC1 was selected on the FSHR ICL3, it was crucial to confirm its binding to the full-length receptor in native conditions. Here, the interaction between iPRC1 and the human FSHR was assayed by a BRET method using the VHH C-terminally fused to the Venus BRET acceptor, and the FSHR or the arginine-vasopressin 2 receptor (AVPR2) fused to the RLuc8 BRET donor. With increasing amounts of transfected VHH-Venus plasmids (Figure 3A), a dose-dependent and much higher binding of iPRC1 to FSHR than to AVPR2 was observed, even though a weak binding signal was detected with AVPR2. There was no impact of the presence of the ligand on iPRC1 binding, which indicates that iPRC1 has no preference for inactive or ligand-induced active conformations of the FSHR. The maximum binding estimated from fitting the data to specific binding curves with Hill slope showed a statistically significant difference for FSHR, when compared to AVPR2 (Figure 3B).

**Figure 3:**
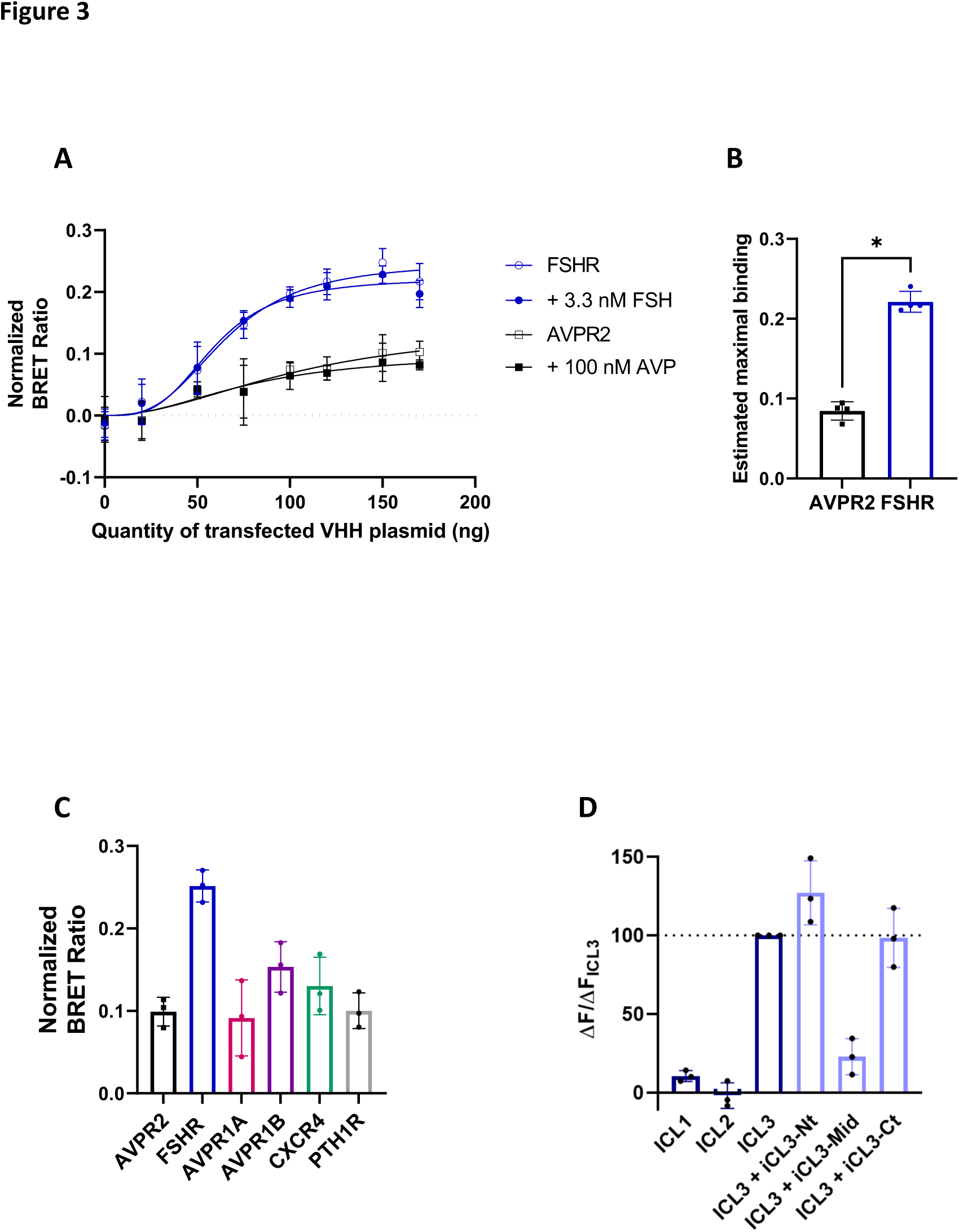
iPRC1 associates to the FSHR, whether FSH is present or not. **(A)** Bioluminescence resonance energy transfer (BRET) binding assay between FSHR-RLuc8 and increasing amounts of transfected iPRC1-Venus plasmid. Experiments were done in the absence of ligand or in the presence of 3.3 nM FSH or 100 nM AVP, as indicated. BRET ratios (at 2 min following FSH addition) were normalized to the BRET ratios obtained in the presence of the irrelevant T31-Venus intra-VHH. Data are represented as mean ± SD, and the curves correspond to fitted data to specific binding curves with Hill slope, N=4. **(B)** Comparison of the estimated maximum binding obtained for iPRC1 to the FSHR to the estimated maximum binding values obtained for iPRC1 association to AVPR2. Data are represented as mean ± SD. Statistical significance was assessed by unpaired Mann-Whitney test, N = 4, * *p* < 0.05. **(C)** BRET binding assay between FSHR-RLuc8, AVPR2-RLuc8, AVPR1A-RLuc8, AVPR1B-RLuc8, CXCR4-RLuc8 or PTH1R-RLuc8 and a fixed amount of iPRC1-Venus plasmid. HEK293A cells were stimulated with either 3.3 nM FSH, 100 nM AVP, 250 nM SDF1α or 10 nM PTH according to the receptor. BRET ratios after 1 min of stimulation were normalized to the BRET ratios obtained in the presence of the irrelevant T31-Venus intra-VHH. Data are represented as mean ± SD, N=3. **(D)** iPRC1 and ICL1, ICL2 or ICL3 peptides interaction and iPRC1-ICL3 interaction using competing peptides ICL3-Nt, ICL3-Mid and ICL3-Ct by HTRF. Data are normalized to background signal, expressed as the percentage of iPRC1-ICL3 interaction signal and represented as mean ± SD, N=3.

The interaction of iPRC1 with other class A GPCRs (AVPR1A, AVPR1B or CXCR4) or a class B GPCR (PTH1R) was also evaluated with a single high quantity of intra-VHH (Figure 3C). All these GPCRs showed an interaction signal close to AVPR2, thus highlighting the specificity of iPRC1 for FSHR. Interestingly, PTH1R contains a BBxxB basic motif similar to the FSHR. This motif that has been reported to be important in Gs coupling to the FSHR (35). Hence, the possibility that this motif could be the iPRC1 epitope can be ruled out.

To confirm the interaction between iPRC1 and the FSHR ICL3 *in vitro*, the interaction between FSHR ICL1, ICL2 or ICL3 peptides and purified iPRC1 was assessed by HTRF (Figure 3D). No interaction was visualized between iPRC1 and ICL1 or ICL2, but as expected, a strong signal was observed with ICL3. In order to further identify the iPRC1 epitope, the interaction of iPRC1 with ICL3 was assessed in presence of competing decapeptides covering ICL3 Nt part (ICL3-Nt), middle part (ICL3-Mid) and Ct part (ICL3-Ct) (Figure 3D). A significant inhibition of interaction was obtained with the ICL3-Mid peptide, suggesting that iPRC1 epitope likely encompasses amino acids ^558^NPNIVSSSSD^567^. In the 3D-structure of the activated FSHR (PDB: 8I2G), the NPNIV motif appears as a flexible loop that is constrainted between the extra-membrane helices formed by ^552^IYLTVR^557^ in Nt and by ^563^SSSDTRI^570^ in Ct. Hence, being the most exposed, this pentapeptide is the most likely epitope of iPRC1.

### Functional effect of iPRC1 on ligand-induced G protein-dependent signalling

Knowing that ICL2 and ICL3 are important interfaces between Gα proteins and class A GPCRs (6), we focused on G protein-dependent signalling. In order to predict whether iPRC1 would interfere with the Gs protein/ FSHR interaction, we first performed docking studies of a modelled iPRC1 CDR3 on the ICL3 of human FSHR active or inactive structures. Docking of the iPRC1 CDR3 on the inactive FSHR led to 9 clusters, the best one consisting of 83 models, having HADDOCK score of –89.1 +/− 5.9 (Figure 4A). The HADDOCK score is the weighted sum of energies, with the most negative values representing the most energetically favourable models. Although only the CDR3 of iPRC1 was defined as the active residues in the interaction during the docking process, the models generated also suggested the involvement of CDR1 and CDR2. Visual inspection of models indicated several poses of iPRC1 with varying vertical-horizontal orientations located on the external side of ICL3 (Figure 4A, left, for a focused view, see Supplementary Figure 3). The superimposition of the best models from the different clusters onto the FSH-FSHR-Gs complex indicated that the binding of iPRC1 to ICL3 would cause steric clash with Gαs (Figure 4A, right, for a focused view, see Supplementary Figure 3). The degree of steric clash with Gαs varied according to the orientation of iPRC1, being minimized with the horizontal pose. However, such horizontal poses could incur steric clashes with the cell membrane. Steric hindrance between iPRC1 and Gs, and the cell membrane, was also confirmed with the active FSH-FSHR-Gs complex, as the models generated did not provide good HADDOCK scores: the best cluster consisted of only 17 models with HADDOCK score of –1.6 +/− 25.1. Overall, it is clear from these models that iPRC1 was either targeting intra-membrane portions of the FSHR or was in steric clash with the cell membrane. Since these docking data suggest a competition between G proteins and iPRC1 upon FSH binding, we next investigated whether iPRC1 would impact ligand-induced G protein-dependant second messenger production. The outcome of iPRC1 or T31 intracellular expression was assayed on cAMP production that results from G protein recruitment upon receptor activation, using HTRF (Figure 4). When compared to the mock-transfected condition (no VHH transfected), iPRC1 induced a decrease of 33% of cAMP accumulation in HEK293A cells transiently co-expressing the FSHR and the intra-VHH, and stimulated for 15 min with 0.3 nM FSH. No effect of the T31 control was observed in the same conditions (Figure 4B). The selectivity of iPRC1 was tested on two other Gs-coupled class A GPCRs, structurally unrelated to the FSHR, the AVPR2 and the beta 2 adrenergic receptor (ADRB2). The AVP-induced cAMP production in HEK293A cells transiently expressing AVPR2, and stimulated for 15 min with 0.25 nM AVP (Figure 4C), or the ISO-induced cAMP production in HEK293A cells endogenously expressing ADRB2 (Figure 4D) were not altered by the co-expression of T31 as expected, nor of iPRC1. These data show that iPRC1 negatively regulates the FSH-induced Gs protein-dependent signalling, and this effect is specific of the anti-FSHR intra-VHH selected in this study.

**Figure 4:**
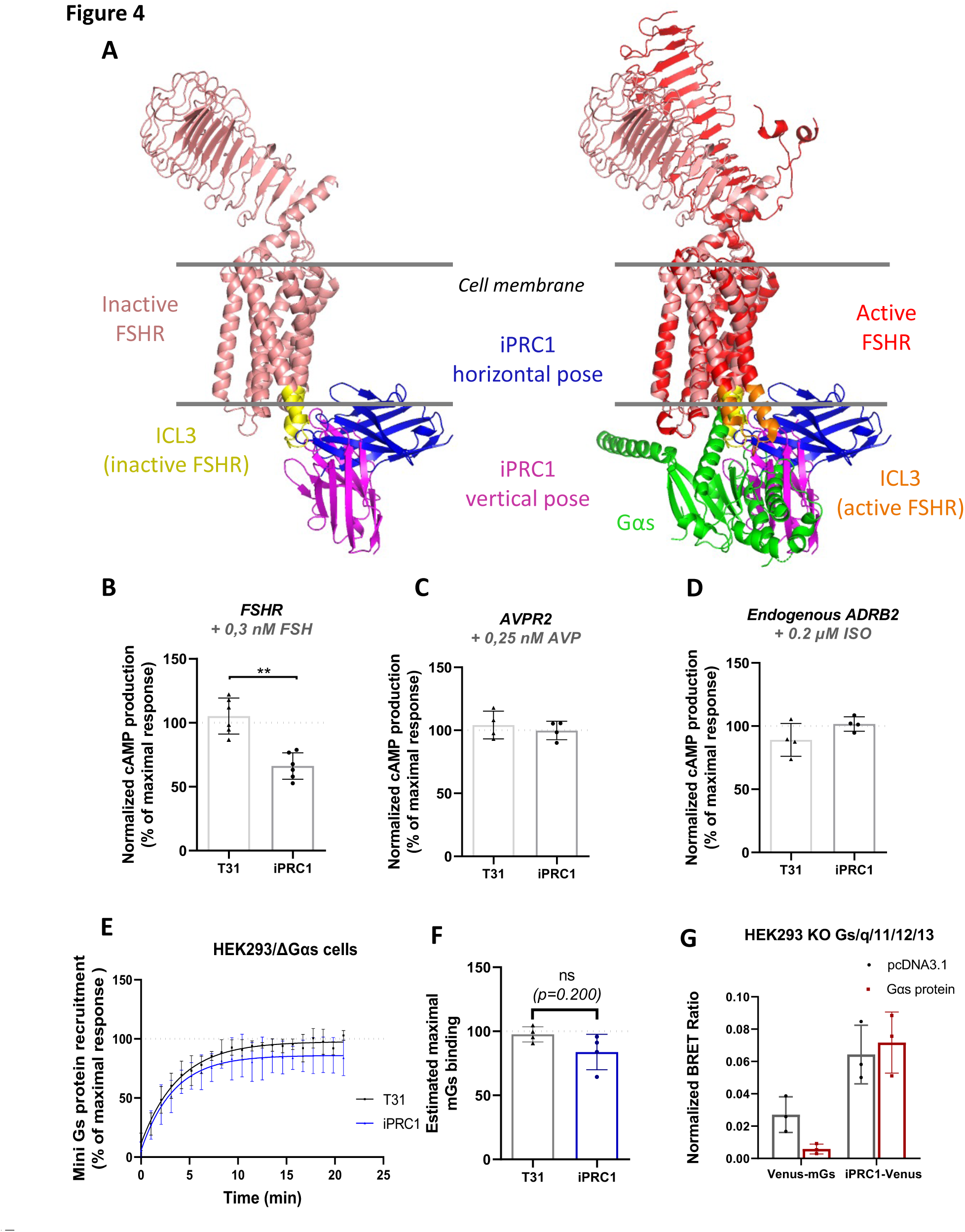
iPRC1 expression affects G protein signalling pathway. **(A)** iPRC1 CDR3 molecular docking to hFSHR ICL3. *Left:* Models of inactive FSHR (PDB: 8I2H) with 2 examples of iPRC1 poses. Right: Models of inactive FSHR (PDB: 8I2H)-iPRC1 superimposed onto FSHR-FSH-Gαs (PDB: 8I2G) (for simplicity, FSH, Gβ, Gγ and Nb35 are not shown). Homogeneous time-resolved fluorescence (HTRF) cAMP accumulation assay in HEK293A cells transiently expressing the AVPR2 **(B)** or FSHR **(C)** or endogenously expressing ADRB2 **(D)**, and transiently expressing either T31 or iPRC1, stimulated with 0.25 nM AVP (N=4), 0.3 nM FSH (N=6) or 0.2 µM ISO (N=4) for 15 min. For each receptor, data were normalized to the cAMP response to Fsk and represented as the percentage of the cAMP response to FSH in the absence of transfected VHH. Statistical significance between the mean ± SD was assessed by unpaired Mann-Whitney test, N = 6, ** *p* < 0.01. **(E)** Bioluminescence resonance energy transfer (BRET) assay measuring the recruitment of NES-Venus-mGs to FSHR-RLuc8, in presence of either T31 or iPRC1, in response to cells stimulation with 33 nM FSH. BRET ratios are expressed as percentage of the maximal FSH-response in the absence of VHH. Data are represented as mean ± SD, and curves correspond to fitted data to one-phase association curves, N = 4. **(F)** Comparison of the estimated maximum binding of Mini Gs proteins to the FSHR in presence of T31 or iPRC1. Data are represented as mean ± SD. Statistical significance was assessed by unpaired Mann-Whitney test, N = 4. **(G)** Interaction inhibition assay in HEK293 cells depleted of Gs/q/11/12/13 transiently transfected with FSHR-RLuc8. iPRC1-Venus and Venus-mGs were transiently expressed in absence (pcDNA3.1) or in presence of overexpressed Gαs protein. Data are represented as mean ± SD (N = 3).

As suggested by the docking data, the reduced cAMP production in response to FSH might be due to iPRC1 partially preventing Gs protein interaction with the receptor. To verify this hypothesis, we assessed the recruitment of Mini Gs to the receptor in absence or in presence of T31 or iPRC1 by BRET (Figure 4E). This experiment was performed in HEK293/ΔGαs to avoid competition with the endogenous untagged Gs proteins. The intracellular expression of iPRC1 slightly decreased Mini Gs/FSH-activated FSHR BRET signal in comparison to T31 or mock conditions, but not significantly, as shown by the comparison of the estimated maximal mGs binding (Figure 4F). These results suggest that iPRC1 does not compete with Gs for interaction with FSHR. However, to confirm this point, we investigated whether an excess of Gαs could displace iPRC1-Venus from FSHR-RLuc8 in HEK293 cells depleted of Gs/q/11/12/13. No difference in iPRC1–FSHR interaction was observed. In contrast, as expected, an excess of Gαs strongly displaced Venus-mGs interaction with the FSHR. These results support the hypothesis that iPRC1 and Gαs do not compete for FSHR binding.

## Discussion

In this study, we have selected the first intra-VHH targeting a member of the glycoprotein hormone receptor family, the FSHR, while only few VHHs targeting the intracellular side of GPCRs have been functionally characterized to date. This intra-VHH is directed against the FSHR ICL3 and impairs the cAMP production of the ligand-bound receptor, despite the proper recruitment of Gαs. Thus, iPRC1 behaves as an allosteric modulator of FSHR.

We first showed that iPRC1 was correctly expressed in mammalian cells and did not aggregate, which makes it well suited for experiments in a cellular context. iPRC1 specifically recognized the FSHR, when compared to other GPCR, Gs-coupled or not. However, iPRC1 specificity towards the two other glycoprotein hormone receptors, LHCGR and TSHR, which share high sequence similarity with the FSHR, still remains to be reported. Importantly, iPRC1 recognised the receptor whether it is FSH-stimulated or not. Hence, iPRC1 is not specific of an inactive or an FSH-activated conformation of the FSHR.

G proteins are recruited to the FSHR notably through the ERW motif present on ICL2, but also through a basic motif on ICL3 (21) (Supplementary Figure 4). We showed that iPRC1 does not bind the PTH1R, a GPCR that contains a basic motif similar to the FSHR’s. Even though these data suggest that this motif is not part of iPRC1 epitope, iPRC1 might interfere with G protein interaction with the FSHR through steric hindrance. This hypothesis is supported by our docking model that overall suggests direct steric clashes with Gαs. Consistently, iPRC1 specifically reduced the FSH-induced cAMP production in HEK293 cells transiently expressing the FSHR, but not the AVPR2 or the ADRB2 in the presence of their respective ligands.

From our results, we anticipated a competition between iPRC1 and G proteins for interaction with the receptor. Surprisingly, when the recruitment of mini Gs to the FSHR was assayed in HEK293 cells depleted of Gαs protein, there was no significant impact of the expression of iPRC1. The affinity of iPRC1 for the FSHR probably being weaker than the affinity of G proteins for the receptor, the binding equilibrium might be in favour of the G proteins. To further investigate if iPRC1 and full Gαs protein are in competition for FSHR recruitment, iPRC1 interaction to the receptor was assessed in HEK293 cells depleted for Gs/q/11/12/13 proteins, in presence of overexpressed Gαs. The latter was not able to displace iPRC1, while it strongly displaced mGs binding, confirming that iPRC1 and Gαs are likely able to bind FSHR simultaneously.

ICL3 is an allosteric site of major interest to modulate GPCR activity and downstream signalling (36). This view probably applies to the FSHR since the recently solved FSHR cryo-EM structure reveals a 14,9 Å movement of the N-term portion of TM6 helix, suggesting the major importance of ICL3 in FSHR activation (16). Therefore, by targeting ICL3, iPRC1 might affect FSHR helix conformational transitions, hence activation. According to the docking studies, an interesting model for the action of iPRC1 would be its binding to the external side of ICL3 in a horizontal pose. Whilst steric clashes with Gαs is minimized, the presence of the iPRC1 however limits the shift of the TM6 to allow for the full recruitment of Gαs since it would be blocked by the cell membrane. This hypothesis is consistent with the binding data suggesting that iPRC1 is able to interact with both active and inactive conformation of FSHR. The Gs protein could also be recruited to the FSHR, but iPRC1 would imply a reorientation that might not be optimal for its activation, thus explaining the reduction of cAMP production. Interestingly, activation of Gαs is accompanied by a prominent shift of helix 5 away from its catalytic domain (6), a movement that may be hindered by the presence of iPRC1. Furthermore, we show that the epitope of iPRC1 overlaps amino acids N^558^ to D^567^ of FSHR ICL3. Yet, according to the recently described structure of the active Gαs-coupled FSHR stabilized with compound 716340, where the ICL3 structure is resolved, amino acid S^566^ engages an interaction with Y^358^ of the Gαs protein (16) (PDB 8I2G). This interaction would be disrupted by iPRC1, giving a possible explanation for partial activation of the Gαs protein.

The ICL3 length exhibits striking diversity among GPCRs, ranging from 10 to 240 amino-acids, and it has been recently proposed that this length might explain G protein selectivity (36), the shorter the loop, the lower the selectivity. In this view, when bound to the FSHR, whose ICL3 is only 23 amino acids long (against 47 and 55 amino acids for AVPR2 and ADRB2 respectively), iPRC1 might promote coupling to other G proteins.

ICL3 being also involved in the interaction of the receptor with β-arrestins, it would be of interest to study the impact of iPRC1 on β-arrestin-dependent signalling, for example through direct β-arrestin recruitment.

Since it does not preferentially interact with an inactive or active conformation of the FSHR, this intra-VHH is also a valuable tool to track FSHR trafficking in gonadal cells, which has never been done before. In addition, an intra-VHH stabilizing a conformation of specific interest could be used in reverse pharmacology. By stabilizing selective conformations of a GPCR with an intra-VHH, it would be possible to screen and identify extracellular binders that stabilize that same conformation and would be suited for therapeutics purposes (37).

Ligand binding to its cognate GPCR results in a series of conformational changes that propagate through the transmembrane domains, allowing the transmission of an extracellular signal to the intracellular compartments of the cell. Although intra-VHH have been instrumental to provide precious structural data on GPCRs, the extreme complexity of GPCR signalling in correlation with conformational changes has not been unravelled. Even though many published structures of GPCRs are available, domains such as the ICLs are unstructured, thus preventing any prediction of structure/ activity relationships. Therefore, intra-VHHs that bind to ICLs and interfere with GPCR signalling pathways constitute valuable tools to help complete structure/ activity studies performed so far.

## Abbreviations

GPCR: G protein-coupled receptor
FSH: Follicle-stimulating hormone
FSHR: FSH receptor
LHCGR: Luteinizing hormone/choriogonadotropin receptor
TSHR: Thyroid-stimulating hormone
BSA: Bovine serum albumin
BRET: Bioluminescence resonance energy transfer
HTRF: Homogeneous time-resolved fluorescence
ICL: intracellular loop
AVP: Arginine-vasopressin
AVPR2: Arginine-vasopressin 2 receptor
ISO: Isoproterenol hydrochloride
ADRB2: β2 adrenergic receptor
Fsk: Forskolin
PBS: Phosphate-buffered saline
RLuc8: luciferase from the sea pansy *Renilla reniformis*
mGs: Mini Gs protein, minimal engineered GTPase domain of the Gα subunit
APC: Allophycocyanine
HEK293: Human Embryonic Kidney 293

## Acknowledgements

The authors would like to acknowledge the Merck company for kindly providing purified recombinant human FSH, Pierre Martineau (Institut de Recherche en Cancérologie de Montpellier, France) for precious technical indications and for kindly providing the pCANTAB 6 phagemid vector and the KM13 helper phage.

## Funding sources and disclosure of conflicts of interest

This work was funded with the support of Institut National de la Recherche Agronomique et de l’Environnement (INRAE), of the MAbImprove Labex (ANR-10-LABX-53) and of Région Centre Val de Loire ARD2020 Biomédicaments SELMAT grant and APR-IR INTACT grant.

PR was funded by a joint fellowship from Région Centre Val de Loire and INRAE PHASE Department. OV was funded by the APR-IR INTACT grant. AV, CaG and VJ was funded by the SELMAT grant of Région Centre Val de Loire ARD2020 Biomédicaments Program. CaG is co-funded by the INTACT grant from Région Centre Val de Loire. TB, ChG, GB and ER are funded by INRAE, FJ-A, NS and PC are funded by the Centre National de la Recherche Scientifique (CNRS).

## SUPPLEMENTARY

**Supplementary figure 1:**
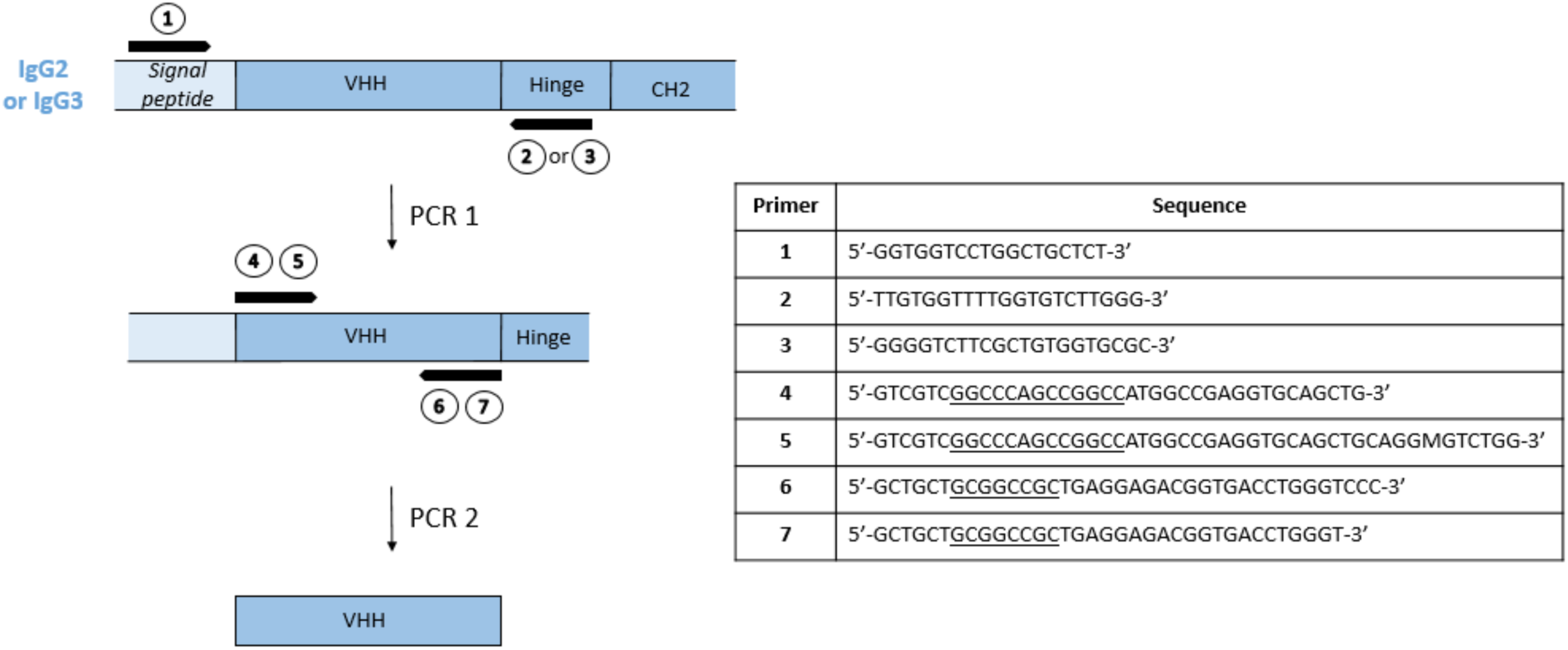
Phage library construction. cDNAs were subjected to a first PCR amplification to discriminate between IgG2 or IgG3 VHH cDNAs and IgG1 VH cDNAs. Two different PCR were performed using **1** as 5’ primer and **2** or **3** specific of the IgG2 and IgG3 hinge regions respectively, as 3’ primers. Four nested PCR were made on each first PCR product using primers encompassing the VHH coding sequence: **4** or **5** from the framework 1 domain as primer 5’, and **6** or **7** from the framework 4 domain as 3’ primer, including the restriction enzyme sites SfiI and NotI (underlined), respectively. The eight nested PCR products were pooled and inserted between the SfiI and NotI sites of the pCANTAB 6 phagemid vector (kindly provided by Pierre Martineau).

**Supplementary figure 2:**
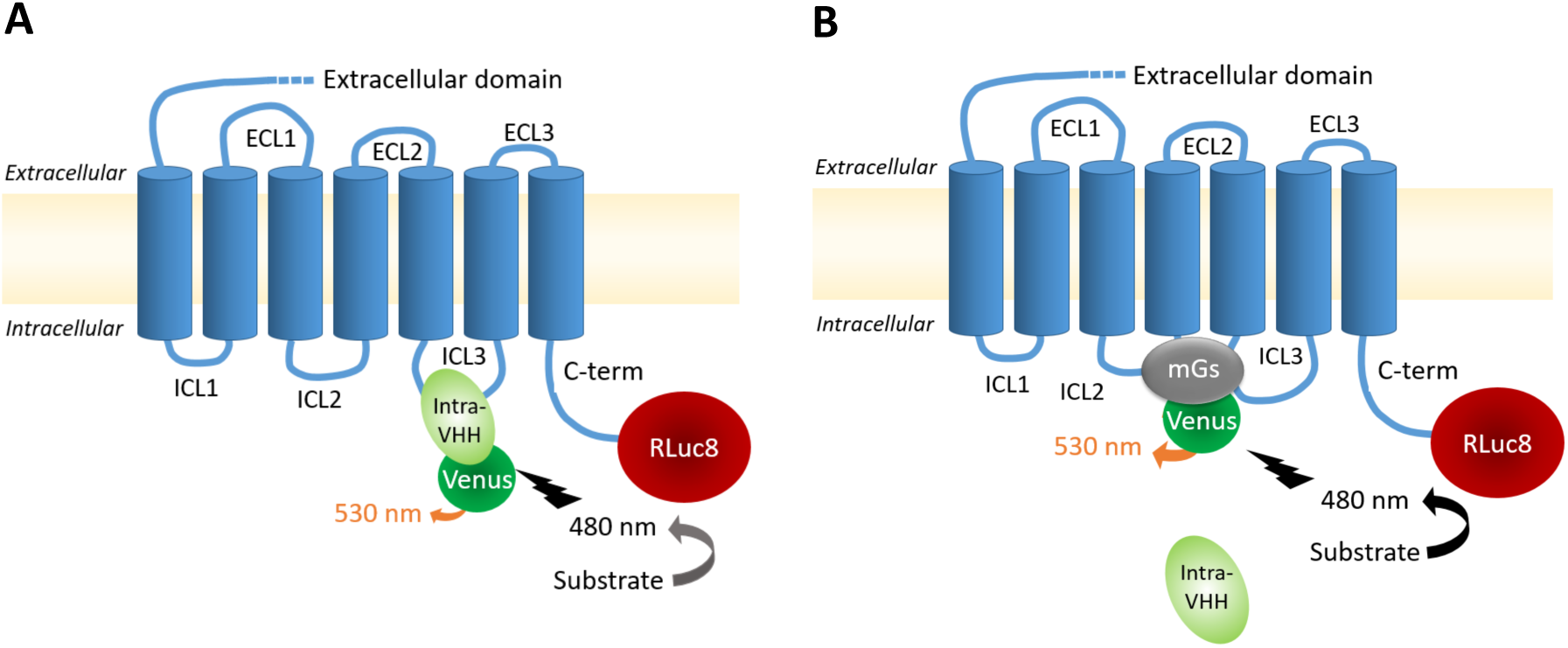
Schematic representation of BRET assays. The RLuc8 fused to FSHR metabolizes its substrate, thus producing a wave at 480 nm, which subsequently excites the Venus fused to the intra-VHH **(A)** or the Mini Gs **(B)** if nearby.

**Supplementary figure 3:**
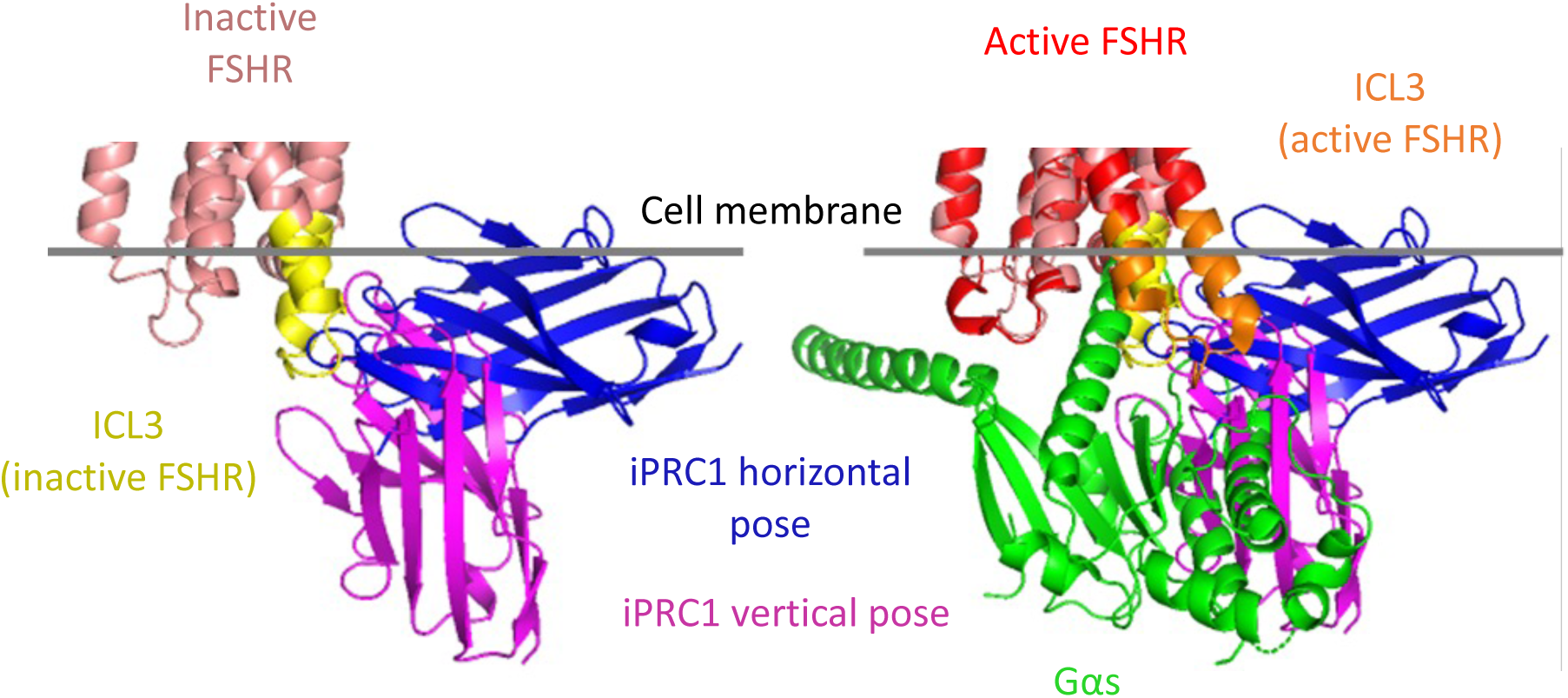
iPRC1 CDR3 molecular docking to the ICL3 at the intracellular side of hFSHR. *Left:* Models of inactive FSHR (PDB: 8I2H) with 2 examples of iPRC1 poses. *Right:* Models of inactive FSHR (PDB: 8I2H)-iPRC1 superimposed onto FSHR-FSH-Gαs (PDB: 8I2G) (for simplicity, Gβ, Gγ and Nb35 are not shown).

**Supplementary figure 4:**
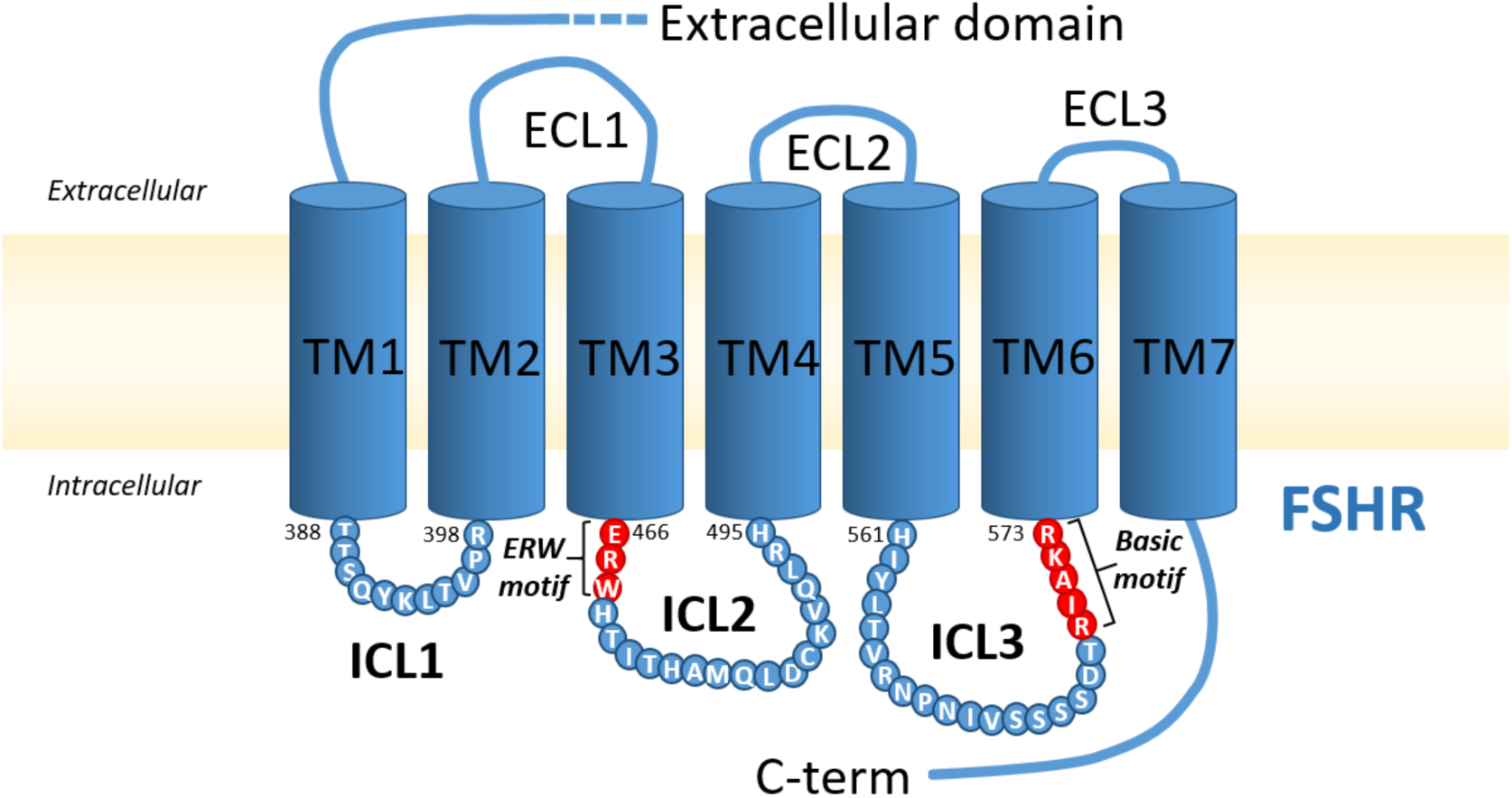
Sites of G protein binding on the FSHR. G proteins are notably recruited to FSHR through two motifs: the ERW motif located on the ICL2, and basic motif (BBXXB) located on the ICL3. *TM: transmembrane domain; ECL: extracellular loop; ICL: intracellular loop*

**Supplementary Table 1:**
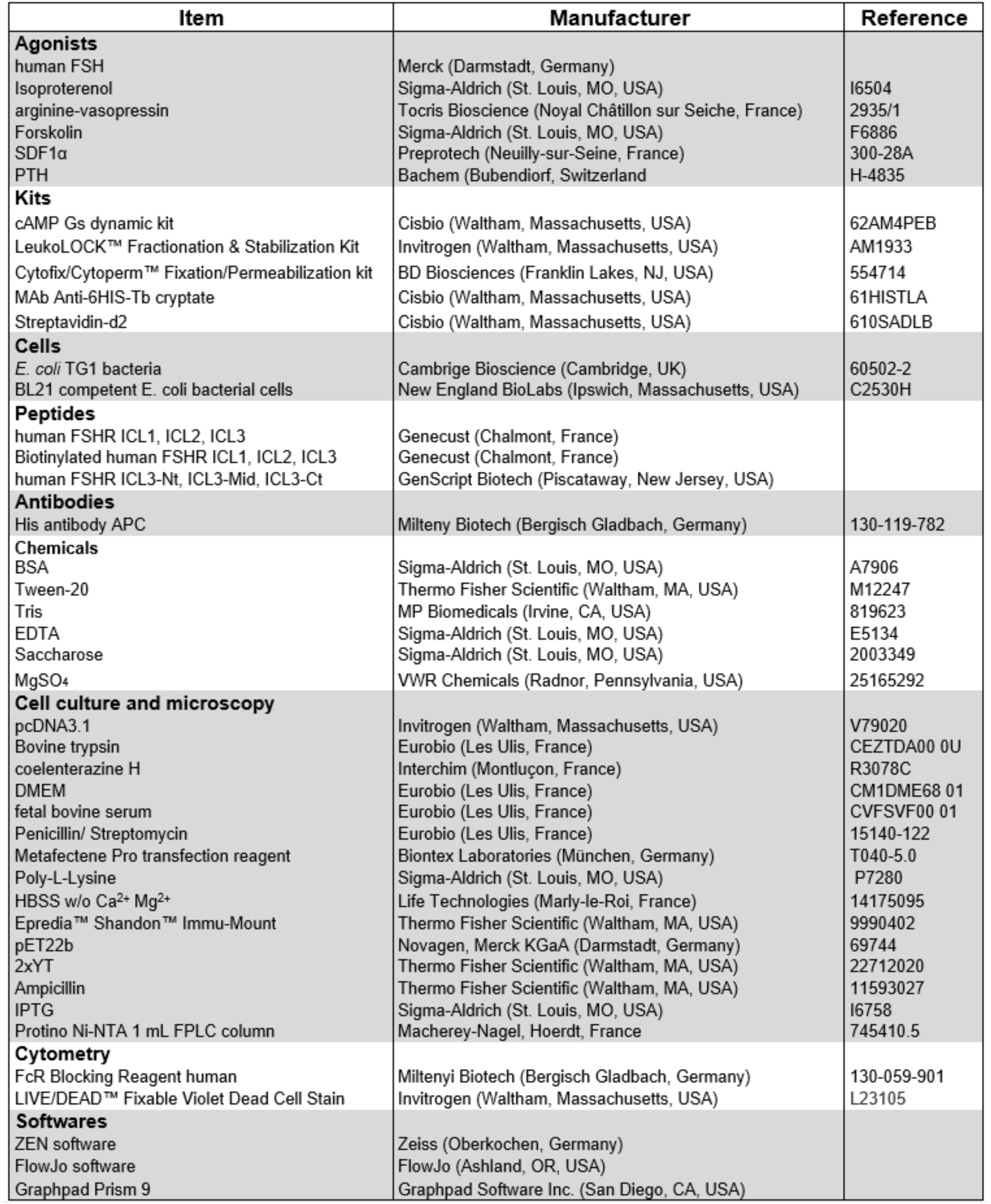
References of the materials.

## Notes

### Competing Interest Statement

The authors have declared no competing interest.

### Summary of Updates

In this revised version, the iPRC1 epitope has been mapped more precisely and its specificity for the FSHR has been challenged with 5 unrelated GPCRs. In addition, our hypothesis that it does not disrupt the G-alpha S recruitment has been demonstrated by competition experiments in G protein-depleted HEK293 cells.

